# Coformulation with tattoo ink for immunological assessment of vaccine immunogenicity in the draining lymph node

**DOI:** 10.1101/2020.08.27.270975

**Authors:** Isaac M Barber-Axthelm, Hannah G Kelly, Robyn Esterbauer, Kathleen Wragg, Anne Gibbon, Wen Shi Lee, Adam K Wheatley, Stephen J Kent, Hyon-Xhi Tan, Jennifer A Juno

## Abstract

Characterisation of germinal centre B and T cell responses yields critical insights into vaccine immunogenicity. Non-human primates are a key pre-clinical animal model for human vaccine development, allowing both lymph node and circulating immune responses to be longitudinally sampled for correlates of vaccine efficacy. However, patterns of vaccine antigen drainage via the lymphatics after intramuscular immunisation can be stochastic, driving uneven deposition between lymphoid sites, and between individual lymph nodes within larger clusters. In order to improve the accurate isolation of antigen-exposed lymph nodes during biopsies and necropsies, we developed and validated a method for co-formulating candidate vaccines with tattoo ink, which allows for direct visual identification of vaccine-draining lymph nodes and evaluation of relevant antigen-specific B and T cell responses by flow cytometry. This approach improves the assessment of vaccine-induced immunity in highly relevant non-human primate models.

## Introduction

Peripheral lymphoid tissues, including lymph nodes (LN), tonsils and mucosal associated lymphoid tissues are critical sites for the generation of adaptive immunity and immunological memory. After intramuscular (IM) administration, vaccine antigens drain via the lymphatics to be concentrated and retained within the LN, where they are subject to immune surveillance. Antibodies are a key protective correlate for most human vaccines, with high-affinity variants generated within germinal centres (GC) via tightly regulated interactions of antigen, GC B (B_GC_) cells, and T follicular helper (T_fh_) cells (1). Efficient generation of GC by immunisation is therefore a key determinant of vaccine success or failure, controlling the kinetics, magnitude and quality of the resultant serological response (2, 3). Direct characterisation of antigen-specific B_GC_ or T_fh_ cells can provide important insights into vaccine immunogenicity and the biogenesis of protective immune responses. However, interrogation of LN B_GC_ and T_fh_ cells in humans is challenging, requiring invasive surgical excision or the collection of a small number of cells by fine needle aspirates (FNA) (4, 5). In contrast, pre-clinical animal models such as non-human primates (NHPs) offer the opportunity to collect longitudinal LN and peripheral blood samples during vaccine studies (6, 7). A factor critical to the detection of these immune responses is the accurate sampling of LNs that drain the injection site, which can be technically challenging due to the sporadic route of antigen trafficking in vivo.

There are multiple factors that can confound accurate sampling of vaccine draining LNs. In humans, IM vaccination into the deltoids sees antigen drain predominately to the axillary LN, with vaccine responses largely restricted to draining LNs in anatomic proximity to the injection site (8–12). Vaccination in the quadriceps is expected to drain predominately to the deep inguinal lymph nodes, which subsequently drains to the external iliac LN. (9, 13, 14). In some individuals, lymphatic drainage from the proximal pelvic limb musculature may bypass the ipsilateral inguinal LNs, and drain directly into the iliac LNs (14, 15). While lymphatic drainage patterns of the thoracic limb are conserved in rodents and NHPs (2, 16, 17), pelvic limb lymphatics predominately drain to the iliac LN, with inconsistent drainage to the ipsilateral inguinal LN (2, 17, 18). In larger species, LNs are grouped in clusters within individual anatomic sites with dissimilar amounts of antigen deposition, further complicating accurate sampling of the vaccine draining LN (2, 17, 19, 20). We and others observed substantial variability in vaccine induced responses when random LN in the draining region are sampled in NHPs (19) and humans, likely in part due to not sampling the responding LN. The ability to directly track antigen drainage following vaccination would substantially improve the accuracy of LN biopsies and assessment of GC immune responses, particularly in large animals such as NHPs. While this can be partially mitigated by substituting subcutaneous (SC) for IM vaccine administration (2), the majority of human vaccines are given IM and it is desirable to maintain comparable delivery in pre-clinical animal models.

Previous studies have used tracking dyes to broadly identify LN drainage patterns in rodent and NHP animal models (2, 16–18, 20–22).Tracking dyes have also been utilised clinically to identify sentinel LNs in cancer patients for biopsy or surgical resection (23–27). However, the potential for mixing vaccine antigens with tracking dyes for long-term demarcation of draining LNs *in vivo* is untested. Here we show that that co-formulating influenza haemagglutinin (HA) or severe acute respiratory syndrome coronavirus 2 (SARS-CoV-2) spike immunogens with adjuvant and tattoo ink allows for the ready visual identification of draining LN without compromising downstream analyses of cellular and humoral immunity. We propose tattoo ink-based vaccine tracking is an effective method for the differentiation of vaccine-draining LNs during extended periods of time post-vaccination, and facilitates a more accurate quantification and phenotypic characterisation of vaccine-specific B and T_fh_ cells in draining LNs, in both murine and NHP animal models.

## Results

### Tattoo ink demarks draining lymph nodes in vivo

Tracking dyes used in vaccinations should (i) visibly stain the draining LN with co-deposition of antigen, and (ii) not affect vaccine immunogenicity or downstream analyses, such as flow cytometric quantification of antigen-specific B_GC_ and T_fh_ populations. We first tested the impact of 3 candidate tracking dyes (Evans blue dye, India ink, and tattoo ink) on cell viability and autofluorescence *in vitro*. Human PBMC cultured in media with 0.05% Evans blue dye for 1 hour resulted in substantial cytotoxicity and loss of lymphocytes (**Fig 1A**). In contrast, culture with 1% India ink or 1% tattoo ink did not affect lymphocyte viability (**Fig 1B**). Despite the short duration of co-culture (1hr), incubation of PBMC with 1% India ink demonstrated alterations in cellular autofluorescence as measured by flow cytometry on channels off the blue, violet and UV lasers (**Fig 1C**). Given tattoo ink demonstrated less autofluorescence (**Fig 1C**), we proceeded to test the utility of tattoo ink co-formulated with vaccine antigens in mice.

**Figure 1 –.**
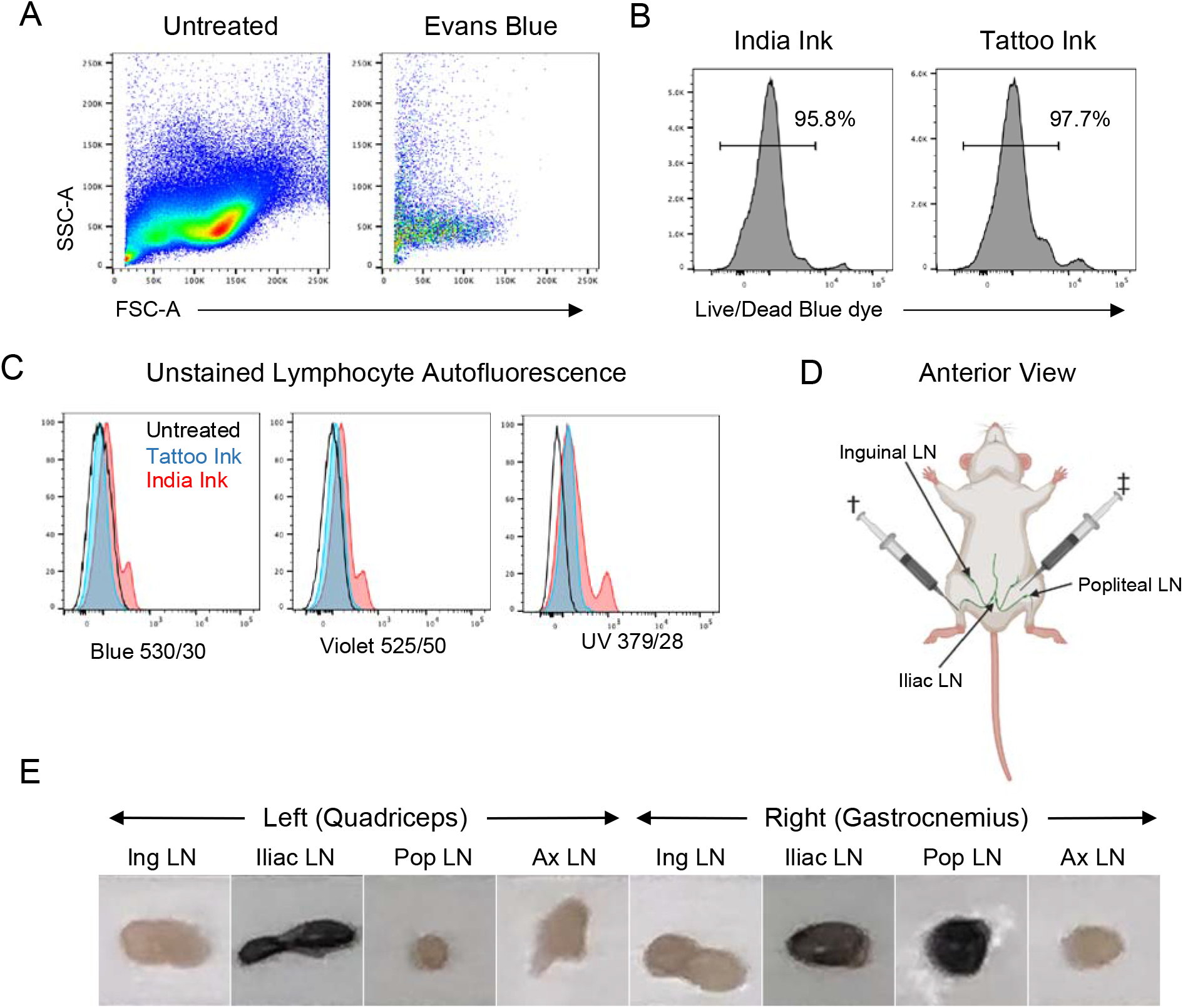
Assessment of candidate dyes for tracking of vaccine-draining LNs. (**A**) Forward and side scatter plots of human PBMC incubated for 1 hour without dye or with 0.05% Evans blue dye. (**B**) Representative viability staining of human PBMC incubated for 1 hour with 1% India ink or 1% tattoo ink. (**C**) Representative histograms of human lymphocyte fluorescence after 1 hour of incubation with no dye (untreated), 1% India ink, or 1% tattoo ink. (**D**) Illustration of vaccine administration sites in the left quadriceps and right gastrocnemius. (**E**) Representative images of draining and non-draining LNs from 1 mouse. †: Right gastrocnemius vaccine; ‡: Left quadriceps vaccine

C57Bl/6J mice were immunised IM in the right gastrocnemius and left quadriceps with A/Puerto Rico/8/1934 haemagglutinin (PR8-HA; 5μg) formulated with Addavax adjuvant and tattoo ink (0.5%). Based on previous studies (16, 18), antigens delivered to the right gastrocnemius will predominately drain to the right popliteal LN and the right iliac LN, with variable drainage to the right inguinal LN (**Fig 1D**). Antigens delivered in the left quadriceps will predominately drain to the ipsilateral iliac LN, with variable drainage to the ipsilateral inguinal LN (**Fig 1D**). Lymph nodes were harvested and assessed visually for ink staining 14 days post-vaccination. On the left quadricep side, non-draining LN (left popliteal and axillary LN) showed no evidence of ink uptake, with the left iliac LN consistently exhibiting ink uptake (**Fig 1E, Table 1**). In contrast, right popliteal and right iliac LNs exhibited obvious ink uptake by eye following right gastrocnemius injection, while absent in the right axillary LN (**Fig 1E, Table 1**). These observations are consistent with previous reports that lymphatics from the pelvic limbs drain into the iliac LNs in mice (16, 18). Dye labelling of inguinal LN was variable, with approximately 20% of the left inguinal LNs, and 60% of the right inguinal LNs labelled (**Fig 1E, Table 1**). This may reflect differences in lymphatic drainage patterns between proximal and distal muscle groups of the pelvic limb. Overall, our results indicate that tattoo ink can label draining LNs when administered in combination with antigen without substantially impacting lymphocyte viability and autofluorescence.

**Table 1:** Frequency of tattoo ink labelling in murine LNs.

|  | Left |  |  |  | Right |  |  |  |
| --- | --- | --- | --- | --- | --- | --- | --- | --- |
|  | Ing. LN | Iliac LN | Pop. LN | Ax. LN | Ing. LN | Iliac LN | Pop. LN | Ax. LN |
| # of LNs with tattoo ink (of n=5 mice) | 1 | 5 | 0 | 0 | 3 | 5 | 5 | 0 |
| Freq. of LN with tattoo ink (%) | 20 | 100 | 0 | 0 | 60 | 100 | 100 | 0 |

Shaded cells indicated lymph nodes that are expected to drain the vaccine sites

### Coformulation with tattoo ink does not compromise flow cytometric immune assessment of vaccine immunogenicity

To assess the extent to which ink staining tracks with vaccine-induced GC activity in mice, we quantified both total B_GC_ (B220^+^IgD^-^GL7^+^CD38^dim^) or HA-specific B_GC_ cell frequency (28) in LNs with and without gross ink uptake (**Fig 2A**). Expanded frequencies of B_GC_ cells were observed in ink-dyed draining LN compared to non-draining LN or undyed draining LN (median 24.6% for ink-dyed LN versus 4.8% for non-draining LN and 8.9% for undyed draining LN; **Fig 2A, B**). Similarly, HA-specific B_GC_ cells were substantially enriched in draining ink-dyed LN (median 10.2% of total B_GC_) compared to non-draining (median 0.0% of total B_GC_) or undyed draining LN (median 1.1% of total B_GC_) (**Fig 2A, B**). These data indicate that the tattoo ink vaccine formulation does not hinder the identification of total or antigen-specific B_GC_ by flow cytometry, and that ink-dyed draining LN are more likely to contain vaccine responses than undyed LN.

**Figure 2.**
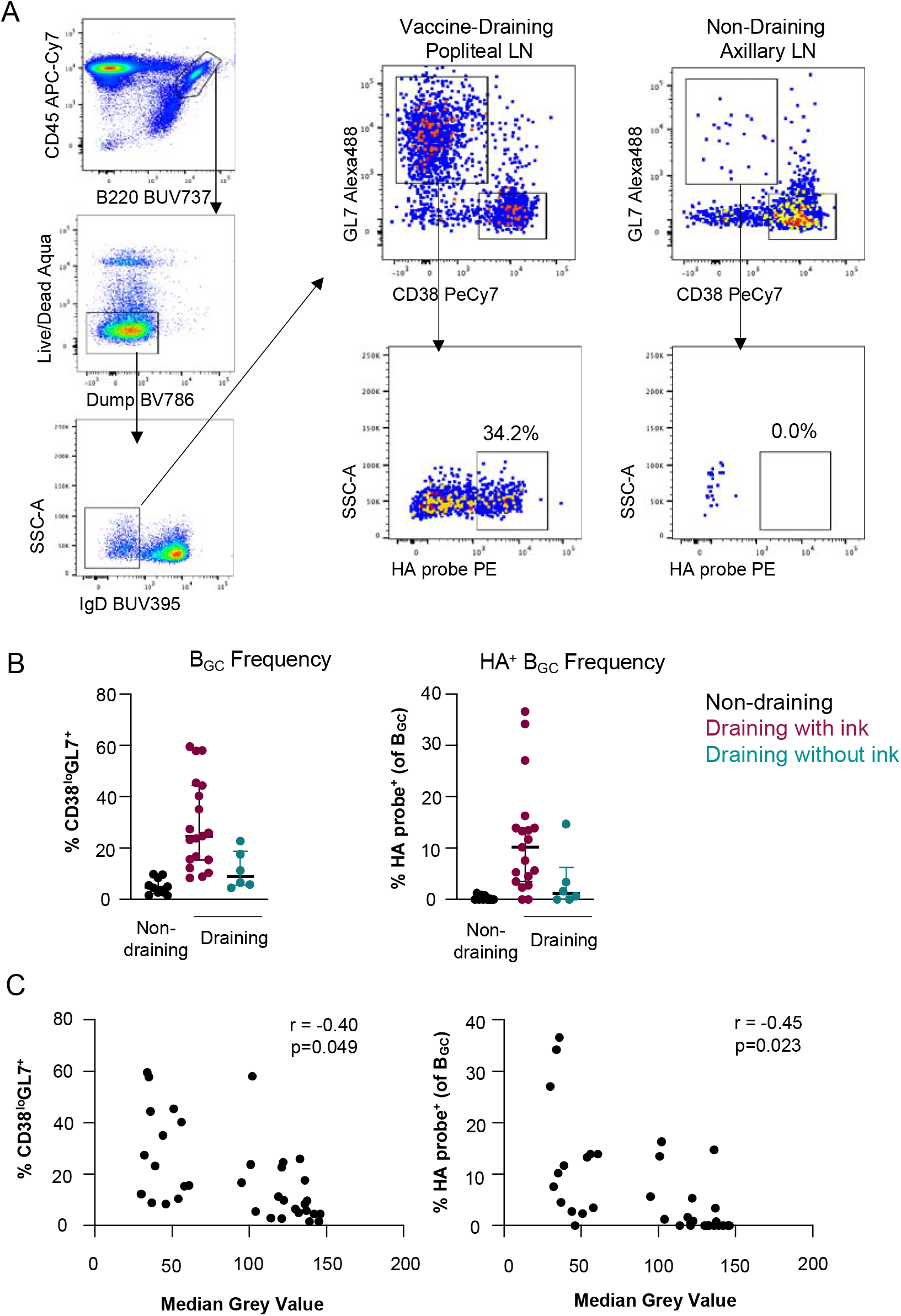
Enrichment of influenza HA-specific germinal centre responses in ink-stained lymph nodes. (**A**) Gating strategy for the identification of HA-specific BGC. Representative plots are shown for a vaccine-draining LN (right popliteal) and a non-draining LN (axillary). (**B**) Quantification of BGC and HA-specific BGC in non-draining LN (axillary and left popliteal), ink-stained draining LN (right popliteal, left/right iliac or left/right inguinal), and non-ink stained draining LN. Each point represents an individual LN collected from a total of n=5 animals vaccinated with HA/tattoo ink mix. Lines and error bars indicate median and IQR. (**C**) Median grey value for each LN was determined by median intensity analysis of 8bit images of the LNs in FIJI/ImageJ. Plots show the correlation between BGC or HA-specific BGC frequency and LN grey value for all draining LN (n=5 animals). Statistics assessed by Spearman correlation.

To further evaluate the utility of tattoo ink in labelling vaccine draining LNs, we performed a quantitative evaluation of the amount of tattoo ink on the surface of draining and non-draining LNs and evaluated the correlation to the frequency of B_GC_ cells and HA-specific B_GC_ cells. Quantitation of tattoo ink deposition in LNs was accomplished by identifying the median intensity value of the surface of the LN, which were taken from gross LN images that were converted from RGB to 8bit images. The median values were subsequently plotted on an 8bit grey scale (0–255), where lower median values correspond to darker LNs, which indicates more tattoo ink uptake. Draining LNs with visible tattoo ink clustered towards lower median grey values, while non-draining LNs and draining LNs without gross tattoo ink deposition clustered towards higher median grey values, indicating that median grey values is suitable quantitative approximate of tattoo ink deposition in LNs. Draining LN median grey value significantly correlated with both total and HA-specific B_GC_ (p=0.049 and 0.023, respectively; **Fig 2C**), suggesting that the degree of ink accumulation mirrors antigen load in the LN following vaccination.

### Differentiation of stochastic antigen drainage patterns in pigtail macaques by tattoo ink

While mice generally have only 1-2 LNs at each lymphoid site (16, 18), primates and humans commonly exhibit chains or clusters of 2-14 LNs (9, 13, 17, 20). This creates additional complexities when evaluating the adaptive immune response, as vaccine antigen may drain to a small subset of the LNs at a given site. We tested if sampling accuracy, and the characterisation of vaccine-elicited immune responses ex vivo, could be improved using tattoo ink to label vaccine-draining LNs in NHPs.

Pigtail macaques (*Macaca nemastrina*) were immunised in the right quadriceps with SARS-CoV-2 spike (100μg) formulated with monophosphoryl lipid A (MPLA) liposomal adjuvant (29), and were boosted IM in the right and left quadriceps with SARS-CoV-2 spike (100μg) formulated with MPLA and tattoo ink (1.0%). Administration at these sites was expected to drain primarily to the left and right iliac LNs, with inconsistent drainage to the left and right inguinal LNs (**Fig 3A**) (2, 17). Animals were additionally immunised in the right and left deltoids with human immunodeficiency virus-1 (HIV-1) fixed trimeric envelope protein (SOSIP) vaccines (100μg) formulated with MPLA and 1.0% tattoo ink (ink in the right deltoid only), with expected drainage to the axillary LNs (**Fig 3A**) (2, 17). Animals were humanely euthanized 13-14 days after the second immunisation, and necropsies were performed to evaluate draining and nondraining LNs for the presence of tattoo ink.

**Figure 3 –.**
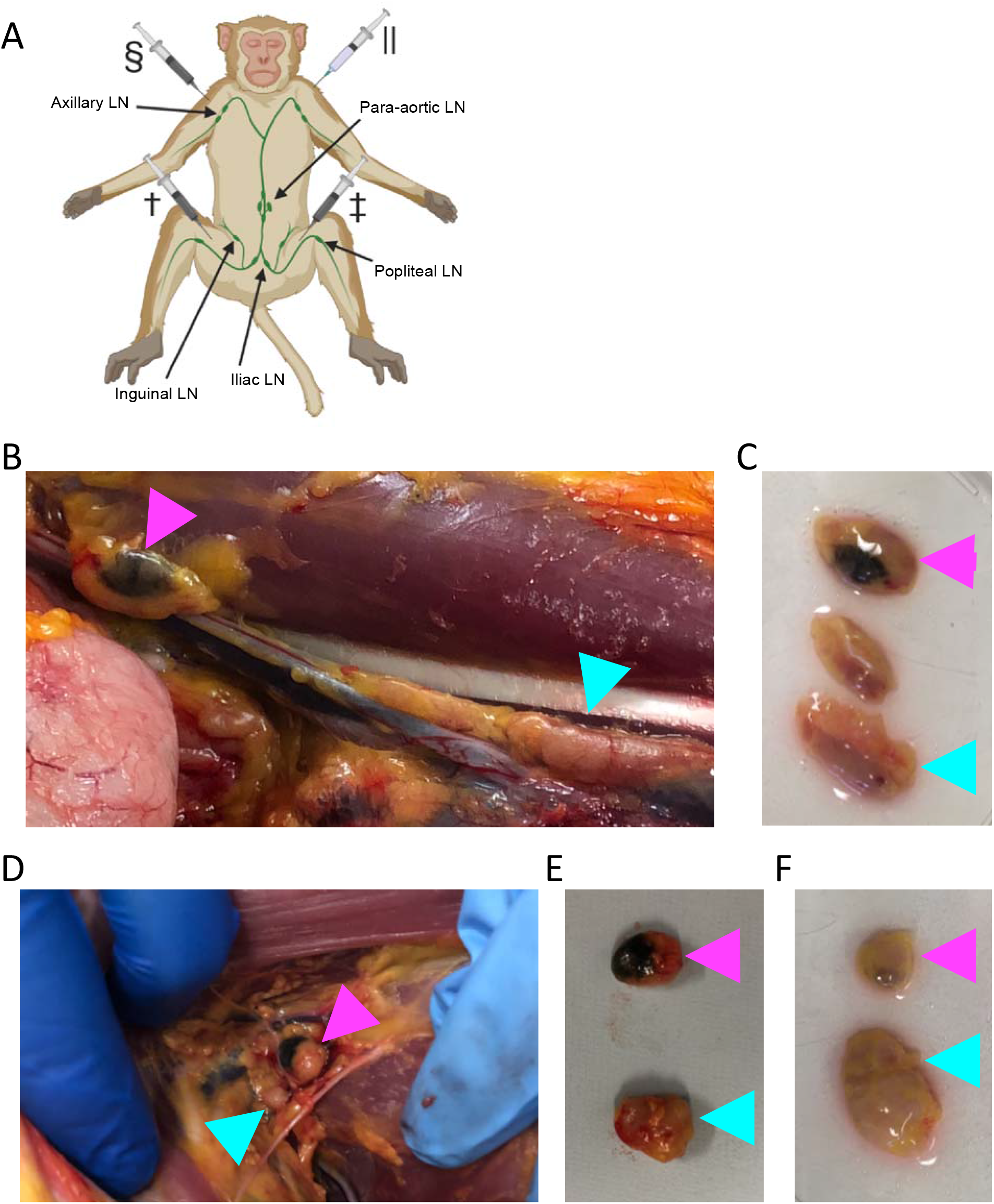
Tattoo ink as a LN tracking dye in non-human primates. (**A**) Illustration of vaccine sites, as well as the associated LNs for intramuscular vaccination in the left and right quadriceps. (**B and C**) NM269 R. iliac LNs in situ (**B**) and after collection (**C**), illustrating the presence of LNs with tattoo ink (magenta arrowhead) and without tattoo ink (cyan arrowhead) within the same chain. (**D and E**) NM269 R. axillary LNs in situ (**D**) and after collection (**E**), illustrating the presence of LNs with tattoo ink (magenta arrowhead) and without tattoo ink (cyan arrowhead) within the same chain. (**F**) NM269 L. axillary LN after collection, illustrating the presence of LNs with tattoo ink (magenta arrowhead) and without tattoo ink (cyan arrowhead) within the same chain. †: Right quadriceps vaccine; ‡: Left quadriceps vaccine; §: Right deltoid vaccine; ∥: Left deltoid vaccine

Among the draining LNs, tattoo ink was visible in at least one LN within the right and left iliac chains in 7 of 8 animals. (**Table 2**). Dye labelling of inguinal LNs was variable, with LNs containing tattoo ink being identified in the left and right inguinal lymphoid sites found in 1 of 8 and 4 of 8 animals, respectively (**Fig 3 B and C, Table 2**). Dye labelling was observed in the right axillary LNs in 8 of 8 animals, while dye labelling in the left axillary LNs was observed in 1 of 8 animals (**Fig 3D-F, Table 2**). Stochastic drainage to the ipsilateral inguinal LN aligns with previous reports (2), and is important to consider as the inguinal LN is a common and readily accessible site for LN sampling via FNA or biopsy. No tattoo ink was grossly visible in any non-draining LN clusters; including the popliteal, para-aortic, mesenteric, mediastinal, tracheobronchial and submandibular LNs (data not shown). Tattoo ink was generally observed in only a single, or limited number of LN recovered from a given lymphoid site (**Fig 3B-F**), suggesting the widely used practice of pooling all LN for immunological analysis could result in significant dilution of vaccine-specific responses and unpredictable effects on reported frequencies of antigen-specific B and T cell responses.

**Table 2:** Frequency of tattoo ink labelling in NHP LNs.

|  | Left |  |  | Right |  |  |
| --- | --- | --- | --- | --- | --- | --- |
|  | Ing. LN | Iliac LN | Ax. LN | Ing. LN | Iliac LN | Ax. LN |
| # of LNs with tattoo ink (of n=8 NHPs) | 1 | 7 | 1 | 4 | 7 | 8 |
| Freq. of LN with tattoo ink (%) | 12.5 | 87.5 | 12.5 | 50 | 87.5 | 100 |

Shaded cells indicated lymph nodes that are expected to drain the vaccine sites

### Ink staining preferentially identifies vaccine-responsive inguinal LNs in non-human primates

Longitudinal studies involving LN biopsies in macaques commonly sample the more readily accessible inguinal LN, to which antigen drainage is highly stochastic (2). Among the 8 animals vaccinated IM into the quadriceps, only 4 exhibited some degree of tattoo ink staining of inguinal LNs (NM11, NM88, NM224, and NM251). Samples from 3 animals (NM11, NM88, and NM224) were available to evaluate antigen specific B_GC_ and T_fh_ frequencies in tattoo ink-containing LNs. Comparison of GC T_fh_ and S-specific B_GC_ populations among ink dyed or undyed inguinal LN, as well as a non-draining LN control (submandibular), in NM11 showed a strong enrichment of vaccine responses and GC activity in the ink stained nodes (**Fig 4A**). In LN samples from NM88, only low levels of S-specific B cells were observed and comparable GC T_fh_ frequencies among both draining (inguinal) and non-draining LN, suggesting that vaccine responses in this animal were restricted to the iliac LN (**Fig 4B**). NM224 was the only animal without ink staining in the iliac LN, with dye accumulation only occurring in the inguinal LN. A comparison of the iliac and inguinal LN confirmed that, similar to NM11, vaccine-induced GC responses were associated with the ink stained inguinal LN (**Fig 4C**). Overall, these results demonstrate the utility of tattoo ink for ex-vivo identification of draining LN enriched for vaccine-induced GC responses.

**Figure 4.**
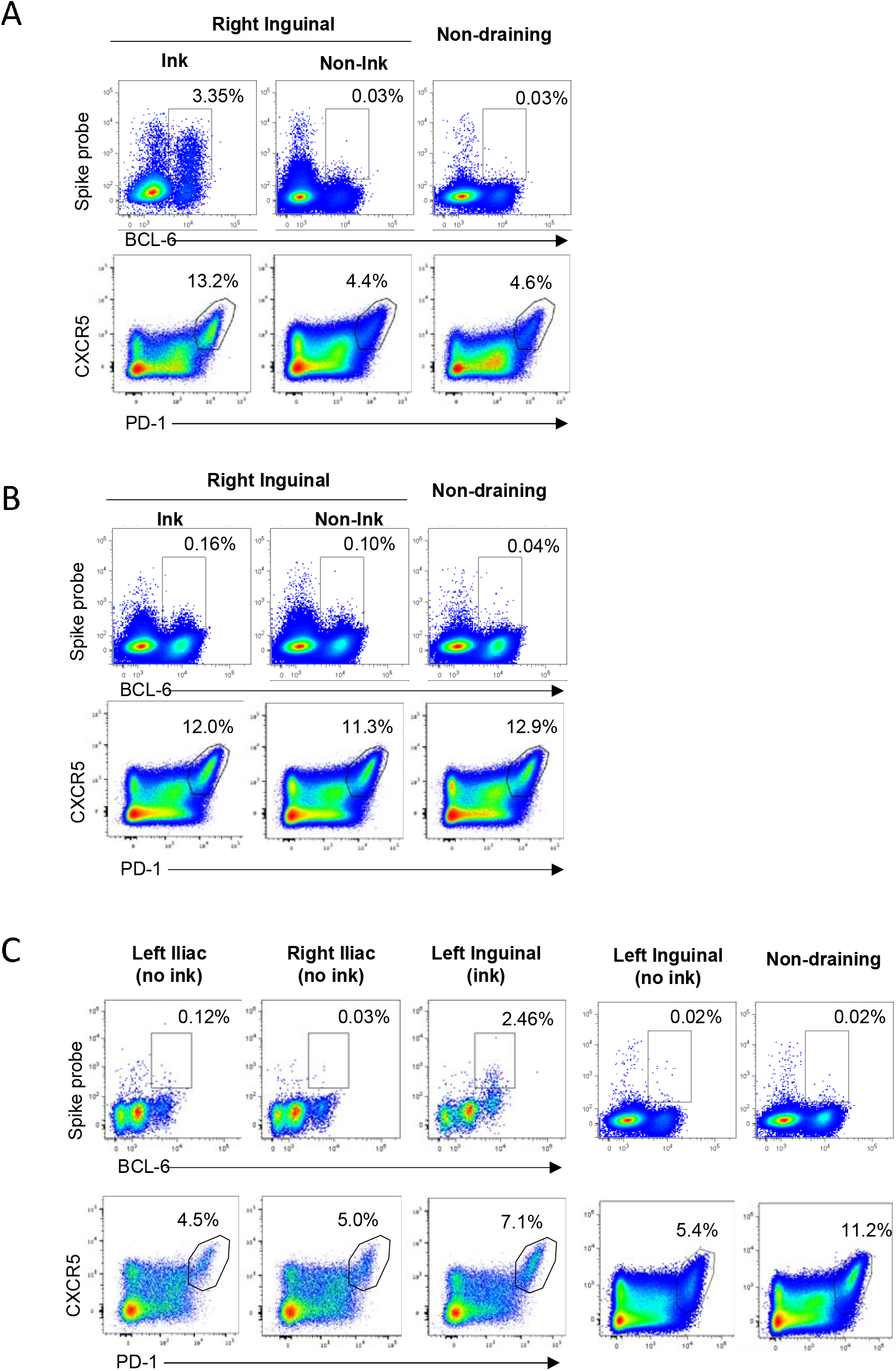
TFH and antigen-specific BGC in stained and unstained LN. (A) Enrichment of GC activity in ink-dyed inguinal LN in NM11 compared to undyed or non-draining (submandibular) LN. (**B**) Lack of vaccine-induced GC activity in any draining inguinal LN in NM88. (**C**) GC activity in NM224, the only animal that exhibited no vaccine responses in the iliac LN. Enrichment of GC activity was observed only in ink-stained inguinal LN.

## Discussion

Draining LNs are key sites for the induction of adaptive immune responses following immunisation and drive the development of immunological memory. A detailed understanding of complicated LN processes, such as the biogenesis of GCs and the induction of antigen-specific T and B cell responses, is key for rational design of new generation vaccines, particularly for pathogens that have eluded traditional vaccine development efforts like HIV and malaria. Techniques enabling the targeted study of vaccine-draining LN allow for precise analyses of immunity without the confounding inclusion of non-draining LN.

Here we show that tattoo ink is an effective and durable tracking dye that can be formulated into vaccines without compromising flow cytometry-based analysis of B_GC_ and T_fh_ cells. Deposited tattoo ink in LNs was associated with higher frequencies of immunogen specific B_GC_ cells in mice, and immunogen-specific B_GC_ and T_fh_ cells in NHPs. We observed high levels of specificity *in vivo*, with minimal staining observed in non-draining LN at distal sites, and good concordance between staining and immunity within a single site. Precise sampling of vaccine draining LNs, therefore, avoids dilutional effects of sampling non-draining LNs from the same site and improves the accurate quantification of antigen-specific B and T cell responses in immunised animals.

Numerous studies employing a variety of different dyes have tracked lymphatic drainage in the hours or days after administration (2, 16–18, 20, 21, 23–27). However, long term stability *in vivo*, and the influence of dye on downstream immunological analysis is unclear. Prolonged labelling of vaccine draining LNs is a relevant consideration, as it allows serial sampling to evaluate changes in the immune response over weeks to months after initial vaccination. Visible dye staining in rats was reported 10-14 days after intraperitoneal administration of pontamine sky blue dye (22). Tattoo ink, by design, is a stable, relatively inert compound that can persist in LNs for extended periods time. In humans, several case reports have noted tattoo ink being incidentally identified in draining LNs up to 30 years after the tattoo was originally placed (30–32). The longevity of tattoo ink in draining LNs when formulated with vaccines still needs to be determined. However, our data demonstrates that draining LNs containing tattoo ink can be readily identified 2 weeks after administration in both mice and NHPs, with the likelihood of it persisting for considerably longer.

Published methods for identifying vaccine-draining LNs include administration of fluorescently labelled immunogens that can be identified by In Vivo Imaging Systems (IVIS) (2, 33), and administration of Tc^99^ sulfur colloid that can be identified with a gamma probe (34). Vaccines may also be administered in specific anatomical locations, or specific routes, to increase the likelihood of antigen drainage to a specific LNs. For example, injection in the SC flank in mice for selective drainage to the ipsilateral inguinal LN (35, 36), or SC immunisation in the anterior thigh for selective drainage to the inguinal and iliac LNs in NHPs (2). The route of vaccine can impact the associated immune response; for example, reports of SC immunisation eliciting stronger neutralising antibody response compared to IM immunisation in NHPs) (37).

Identification of vaccine draining LNs with tattoo ink provides a simple approach for active LN identification without requiring specialised equipment, while permitting IM vaccination routes of greatest clinical relevance for human vaccines.

One consideration with our proposed method is the potential for both endogenous and exogenous pigments to confound the identification of ink containing lymph nodes. Hemosiderin, and iron-containing haemoglobin breakdown product, can accumulate in draining lymph nodes from congested, haemorrhagic, or inflamed areas (38–40). Similarly, carbon-containing, particulate debris can accumulate in tracheobronchial and mediastinal lymph nodes, following inhalation and drainage from the pulmonary tree (38, 39, 41). Accumulations of any of these pigments can cause lymph node darkening which could be mistaken for tattoo ink accumulation, and confound positive identification of relevant lymph nodes. Such hemosiderin accumulation in draining lymph nodes may have contributed to false identification of NM88 right inguinal LN and NM269 left axillary LN as tattoo ink-containing LNs in the current study. Tattoo ink comprising alternative colours could be explored to address this.

Overall, we find that co-formulating immunogens with commercial tattoo ink does not hinder the adaptive immune response in the draining lymph nodes, nor interfere with flow cytometry-based methods of immune analysis. While the addition of tattoo ink did not impact the reported flow cytometry panels, it remains possible that other fluorophores may be negatively affected and compatibility with bespoke flow cytometry panels should be established in vitro prior to in vivo usage. While this study focussed on protein-based vaccines draining to lymph nodes in the pelvic limb, we show potential utility for fore-limb immunisations draining to the axillary LNs, and this approach could be extended to other tissues sites and other vaccine platforms. Increased accuracy in profiling vaccine immunity in key pre-clinical animal models is essential to guide the rational improvement of vaccines for human use.

## Methods

### Mouse studies

Mouse studies and related experimental procedures were approved by the University of Melbourne Animal Ethics Committee (ID#: 1914874). Female C57BL/6JArc mice (6-8 weeks old) were anesthetized by isoflurane inhalation prior to immunisation. For IM vaccinations, 5μg of PR8-HA protein with Addavax (1:1 ratio; InvivoGen) and 0.5% tattoo ink was injected into the left quadriceps and right gastrocnemius (50 μL per site), using a 29G needle. Fourteen days post-vaccination, the mice were euthanized via CO_2_ asphyxiation, and inguinal, iliac, popliteal, and axillary LNs were collected for further analysis.

### Non-human primate studies

Macaque studies and related experimental procedures were approved by the Monash University Animal Ethics Committee (ID #23997). Pigtail macaques (*Macaca* nemistrina) were housed in the Monash Animal Research Platform (MARP) Gippsland Field Station. Eight male pigtail macaques (*Macaca nemestrina*) (6-15 years old) were initially vaccinated with 100μg of SARS-CoV-2 spike protein, consisting of either the whole spike protein (S) or the receptor binding domain of the spike protein (RBD), with 200μg of Monophosphoryl Lipid A (MPLA) adjuvant IM in the right quadriceps (total volume: 1ml). A booster vaccine of 100μg S or RBD protein, with 200μg of monophosphoryl lipid A (MPLA) and 1% tattoo ink, was administered IM in both quadriceps (500 μL per site) 28 days after the initial vaccine. The macaques were concurrently vaccinated in the right and left deltoids with human immunodeficiency virus-1 (HIV-1) fixed trimeric envelope protein gp140 (SOSIP) vaccines (100μg) (42, 43) formulated with MPLA and 1.0% tattoo ink (right deltoid only) (1ml per site). Twenty-four hours prior to necropsy, the macaques received an intravenous infusion of autologous Vδ2^+^Vγ9^+^ T-cells labelled with CellTrace Blue (Life Technologies). Thirteen to 14 days post-booster vaccination, the macaques were euthanized by barbiturate overdose, and draining and non-draining LNs were collected for further analysis.

### Tracking dyes and cell culture

Fresh or cryopreserved human PBMC’s were isolated from heparinised whole blood by Ficoll gradient (Sigma). Cells were incubated for 1hr (37°C, 5% CO_2_) in RPMI (Life Technologies) supplemented with 10% fetal calf serum (FCS) and 5% penicillin/streptomycin (RF10). Samples were incubated with 0.05% W/V Evans blue dye (Sigma Life Sciences), 1.0% V/V India ink (Higgins waterproof ink), or 1.0% V/V tattoo ink (Mom’s millennium Black Pearl) in 6-well plates at ~1.5-2.0e^6^ cells/ml. Control samples were incubated in RF10 without dye, under identical conditions. Samples were washed 2 times with PBS prior to analysis.

### PR8-HA and SARS-CoV-2 protein synthesis and purification

Full length influenza H1N1 A/Puerto Rico/8/34 hemagglutinin (PR8-HA) (19, 44), SARS-CoV-2 spike protein (S) (45), and the receptor binding domain of the SARS-CoV-2 spike protein (RBD) (45), were prepared as previously described, and used for flow cytometry probes or as immunogens. PR8-HA and SARS-CoV-2 S proteins with C-terminal Avi-tags and His-tags were expressed via transient transfection of Expi293 suspension cultures (Thermo Fisher) and purified by polyhistidine-tag affinity and size exclusion chromatography. For flow cytometry B cell probes, purified PR8-HA and S proteins were biotinylated using BirA biotin-protein ligase (Avidity). Biotinylated PR8-HA and S proteins were fluorescently labelled by the sequential addition of streptavidin-conjugated to phycoerythrin (PE) (Thermo Fisher) prior to use.

### Lymph node sample processing

Murine LNs were processed and analysed individually. Prior to processing, all LNs were photographed and noted for the presence or absence of visible tattoo ink. NHP LNs from each animal and anatomical location were separated based on the visual presence or absence of ink, and processed as a pooled sample. Murine and NHP LNs were dissociated and passed through a 70μm filter to generate single cell suspensions. Murine LN suspensions were freshly stained for antigen-specific B cells. NHP LN suspensions were cryopreserved in 90% FCS/10% DMSO.

### Tattoo ink accumulation

The amount of tattoo ink deposition in murine draining LNs was estimated based on the amount visible tattoo ink on the surface of the LN. Vaccine draining LNs (right and left inguinal LNs, right and left iliac LNs, right popliteal LN) and non-draining LNs (left popliteal LN, right and left axillary LNs) were evaluated 14 days-post IM vaccination with PR8-HA (5μg) protein with Addavax (1:1 ratio; InvivoGen) and 0.5% tattoo ink. Draining and non-draining LNs were photographed for analysis, and the images were analysed using FIJI/ImageJ (46, 47). The photographs were converted to 8bit images, and individual LNs were outlined with the polygonal region of interest tool. The amount of tattoo ink deposition in each individual LN was then estimated by measuring the median intensity value within each region of interest. The median values were subsequently plotted on an 8bit grey scale (0–255), where lower median values correspond to darker LNs, which indicates more tattoo ink uptake.

### Antibodies and flow cytometry

Fresh, single-cell suspensions of individual murine lymph nodes were stained with live/dead Aqua viability dye (Life Technologies), and F_c_ blocked with CD16/CD32 (93, Biolegend). Surface staining was then performed with the following antibodies: GL7 Alexa488 (GL7, Biolegend), CD45 APC-Cy7 (30-F11, BD), F4/80 BV786 (BM8, Biolegend), CD3ε BV786 (145-2C11, Biolegend), CD38 PE-Cy7 (90, Biolegend), IgD BUV395 (11-26c.2a, BD), and B220 BUV737 (RA3-6B2, BD). Biotinylated PR8-HA with Streptavidin-PE (Thermo Fisher) was used to identify PR8-HA specific B_GC_ cells, and streptavidin-BV786 (BD) was included to identify non-specific streptavidin binding to the cell surface. After 30 minutes incubation at 4°C, cells were washed and fixed in BD Cytofix fixation buffer. Samples were acquired on a BD LSR Fortessa using BD FACS Diva.

For NHP B-cell analysis, thawed lymph node single cell suspensions were stained with live/dead Aqua viability dye (Life Technologies) and the following surface antibodies: IgD Alexa488 (polyclonal, Southern Biotech), CD20 APC-Cy7 (2H7, Biolegend), CD14 BV510 (M5E2, Biolegend), CD3 AF700 (SP34-2, BD), CD8α BV510 (RPA-T8, Biolegend), CD16 BV510 (3G8, Biolegend), CD10 BV510 (HI10a, Biolegend), IgG BV786 (G18-145, BD), CD95 BUV737 (DX2, BD), CD4 BV605 (L200, BD) CXCR5 PE-Cy7 (MU5UBEE, eBioscience) and PD-1 BV421 (EH12.2H7, Biolegend). S-specific B cells were identified using biotinylated SARS-CoV-2 S probe conjugated to Streptavidin-PE (Thermo Fisher). Streptavidin-BV510 (BD) was included to identify non-specific streptavidin binding to the cell surface. Cells were washed and permeabilised with Transcription Factor Buffer Set (BD) prior to BCL-6 AF647 (IG191E/A8, Biolegend) and Ki-67 BUV395 (B56, BD) staining. Samples were acquired on a BD LSR Fortessa using BD FACS Diva.

For T cell analysis following stimulation, single cell suspensions were stained with live/dead Aqua viability dye and the following cell surface markers: CD20 BV510 (2H7, BD), CD3 Alexa700 (SP34-2, BD), CD4 BV605 (L200, BD), CD8 (RPA-T8, Biolegend), CXCR5 PE (MU5UBEE, ThermoFisher), PD-1 BV421 (EH12.2H7, Biolegend), CD95 BUV737 (DX2, BD), CD25 APC (BC96, Biolegend), OX-40 BUV395 (L106, BD). After 30 minutes incubation at 4°C, cells were washed and fixed in 1% formaldehyde. Samples were acquired on a BD LSR Fortessa using BD FACS Diva.

### Statistical and data analysis

Data are presented as median and interquartile range. Spearman correlation was used to compare the frequencies of B_GC_ and PR8-HA specific B_GC_ cells with the median grey value of the corresponding LN. All statistical analysis data presentation was performed using GraphPad Prism version 8 (GraphPad Software, La Jolla California USA). Flow cytometry data was analysed in FlowJo v9 or v10.

## Acknowledgements

The authors would like to thank the staff at the Monash Animal Research Platform (MARP) and Gippsland Field Station, including Irwin Ryan, for their assistance with the non-human primate study. We thank Robin Shattock (Imperial College London), Marit Van Gils (Amsterdam Medical Centre) and the EAVI consortium for the provision of HIV immunogens, and Dietmar Katinger and Philipp Mundsperger from Polymun for provision of the MPLA liposome adjuvant. This work was funded by NHMRC program grant (APP1149990), an NHMRC-EU collaborative award (APP1115828), the European Union Horizon 2020 Research and Innovation Programme under grant agreement 681137, the Victorian government and the Australian Medical Research Future Fund (GNT2002073). SJK, AKW and JAJ are funded by NHMRC fellowships.

## References

1. Grant, S. M., M. Lou, L. Yao, R. N. Germain, and A. J. Radtke. 2020. The lymph node at a glance - how spatial organization optimizes the immune response. J Cell Sci 133: jcs241828.

2. Havenar-Daughton, C., D. G. Carnathan, A. V. Boopathy, A. A. Upadhyay, B. Murrell, S. M. Reiss, C. A. Enemuo, E. H. Gebru, Y. Choe, P. Dhadvai, F. Viviano, K. Kaushik, J. N. Bhiman, B. Briney, D. R. Burton, S. E. Bosinger, W. R. Schief, D. J. Irvine, G. Silvestri, and S. Crotty. 2019. Rapid germinal center and antibody responses in non-human primates after a single nanoparticle vaccine immunization. Cell Reports 29: 1756–1766.e8.

3. Havenar-Daughton, C., D. G. Carnathan, A. T. de la Peña, M. Pauthner, B. Briney, S. M. Reiss, J. S. Wood, K. Kaushik, M. J. van Gils, S. L. Rosales, P. van der Woude, M. Locci, K. M. Le, S. W. de Taeye, D. Sok, A. U. R. Mohammed, J. Huang, S. Gumber, A. Garcia, S. P. Kasturi, B. Pulendran, J. P. Moore, R. Ahmed, G. Seumois, D. R. Burton, R. W. Sanders, G. Silvestri, and S. Crotty. 2016. Direct probing of germinal center responses reveals immunological features and bottlenecks for neutralizing antibody responses to HIV Env trimer. Cell Reports 17: 2195–2209.

4. Hey-Nguyen, W. J., Y. Xu, C. F. Pearson, M. Bailey, K. Suzuki, R. Tantau, S. Obeid, B. Milner, A. Field, A. Carr, M. Bloch, D. A. Cooper, A. D. Kelleher, J. J. Zaunders, and K. K. Koelsch. 2017. Quantification of residual germinal center activity and HIV-1 DNA and RNA levels using fine needle biopsies of lymph nodes during antiretroviral therapy. Aids Res Hum Retrov 33: 648–657.

5. Havenar-Daughton, C., I. G. Newton, S. Y. Zare, S. M. Reiss, B. Schwan, M. J. Suh, F. Hasteh, G. Levi, and S. Crotty. 2020. Normal human lymph node T follicular helper cells and germinal center B cells accessed via fine needle aspirations. J Immunol Methods 479: 112746.

6. Dai, K., L. He, S. N. Khan, S. O’Dell, K. McKee, K. Tran, Y. Li, C. Sundling, C. D. Morris, J. R. Mascola, G. B. K. Hedestam, R. T. Wyatt, and J. Zhu. 2015. Rhesus macaque B-cell responses to an HIV-1 trimer vaccine revealed by unbiased longitudinal repertoire analysis. Mbio 6: e01375–15.

7. Xu, Y., C. Fernandez, S. Alcantara, M. Bailey, R. D. Rose, A. D. Kelleher, J. Zaunders, and S. J. Kent. 2013. Serial study of lymph node cell subsets using fine needle aspiration in pigtail macaques. J Immunol Methods 394: 73–83.

8. Liang, F., G. Lindgren, K. J. Sandgren, E. A. Thompson, J. R. Francica, A. Seubert, E. D. Gregorio, S. Barnett, D. T. O’Hagan, N. J. Sullivan, R. A. Koup, R. A. Seder, and K. Loré. 2017. Vaccine priming is restricted to draining lymph nodes and controlled by adjuvant-mediated antigen uptake. Sci Transl Med 9: eaal2094.

9. Andrade, M., and A. Jacomo. 2007. Cancer treatment and research. 55–77.

10. Ma, C.-X., W.-R. Pan, Z.-A. Liu, F.-Q. Zeng, Z.-Q. Qiu, and M.-Y. Liu. 2019. Deep lymphatic anatomy of the upper limb: an anatomical study and clinical implications. Ann Anat - Anatomischer Anzeiger 223: 32–42.

11. Shirone, N., T. Shinkai, T. Yamane, F. Uto, H. Yoshimura, H. Tamai, T. Imai, M. Inoue, S. Kitano, K. Kichikawa, and M. Hasegawa. 2012. Axillary lymph node accumulation on FDG-PET/CT after influenza vaccination. Ann Nucl Med 26: 248–252.

12. Coates, E. E., P. J. Costner, M. C. Nason, D. M. Herrin, S. Conant, P. Herscovitch, U. N. Sarwar, L. Holman, J. Mitchell, G. Yamshchikov, R. A. Koup, B. S. Graham, C. M. Millo, J. E. Ledgerwood, and V. 900 S. Team. 2017. Lymph node activation by PET/CT following vaccination with licensed vaccines for human papillomaviruses. Clin Nucl Med 42: 329–334.

13. Pflug, J. J., and J. S. Calnan. 1971. The normal anatomy of the lymphatic system in the human leg. Brit J Surg 58: 925–930.

14. Yamazaki, S., H. Suami, N. Imanishi, S. Aiso, M. Yamada, M. Jinzaki, S. Kuribayashi, D. W. Chang, and K. Kishi. 2013. Three□dimensional demonstration of the lymphatic system in the lower extremities with multi□detector□row computed tomography: A study in a cadaver model. Clin Anat 26: 258–266.

15. Pan, W., F. Zeng, D. Wang, and Z. Qiu. 2017. Perforating and deep lymphatic vessels in the knee region: an anatomical study and clinical implications. Anz J Surg 87: 404–410.

16. Kawashima, Y., M. Sugimura, Y.-C. Hwang, and N. Kudo. 1964. The lymph system in mice. Jpn J Vet Res 12: 69–78.

17. Hayakawa, T. 1994. The lymphatics of Japanese macaque. Anthropol Sci 102: 165–179.

18. Harrell, M. I., B. M. Iritani, and A. Ruddell. 2008. Lymph node mapping in the mouse. J Immunol Methods 332: 170–174.

19. Kelly, H. G., H.-X. Tan, J. A. Juno, R. Esterbauer, Y. Ju, W. Jiang, V. C. Wimmer, B. C. Duckworth, J. R. Groom, F. Caruso, M. Kanekiyo, S. J. Kent, and A. K. Wheatley. 2020. Self-assembling influenza nanoparticle vaccines drive extended germinal center activity and memory B cell maturation. Jci Insight 5: e136653.

20. Gooneratne, B. W. M. 1972. The lymphatic system in rhesus monkeys *(Macaca mulatta)* outlined by lower limb lymphography. Cells Tissues Organs 81: 602–608.

21. Smedley, J., B. Turkbey, M. L. Bernardo, G. Q. D. Prete, J. D. Estes, G. L. Griffiths, H. Kobayashi, P. L. Choyke, J. D. Lifson, and B. F. Keele. 2014. Tracking the luminal exposure and lymphatic drainage pathways of intravaginal and intrarectal inocula used in nonhuman primate models of HIV transmission. Plos One 9: e92830.

22. Tilney, N. L. 1971. Patterns of lymphatic drainage in the adult laboratory rat. J Anat 109: 369–83.

23. Frumovitz, M., E. D. Euscher, M. T. Deavers, P. T. Soliman, K. M. Schmeler, P. T. Ramirez, and C. F. Levenback. 2012. “Triple injection” lymphatic mapping technique to determine if parametrial nodes are the true sentinel lymph nodes in women with cervical cancer. Gynecol Oncol 127: 467–471.

24. Gumus, M., H. Gumus, S. E. Jones, P. A. Jones, A. R. Sever, and J. Weeks. 2013. How long will I be blue? Prolonged skin staining following sentinel lymph node biopsy using intradermal patent blue dye. Breast Care 8: 199–202.

25. Nour, A. 2004. Efficacy of methylene blue dye in localization of sentinel lymph node in breast cancer patients. Breast J 10: 388–391.

26. Zakaria, S., T. L. Hoskin, and A. C. Degnim. 2008. Safety and technical success of methylene blue dye for lymphatic mapping in breast cancer. Am J Surg 196: 228–233.

27. Brahma, B., R. I. Putri, R. Karsono, B. Andinata, W. Gautama, L. Sari, and S. J. Haryono. 2017. The predictive value of methylene blue dye as a single technique in breast cancer sentinel node biopsy: a study from Dharmais Cancer Hospital. World J Surg Oncol 15: 41.

28. Tan, H.-X., S. Jegaskanda, J. A. Juno, R. Esterbauer, J. Wong, H. G. Kelly, Y. Liu, D. Tilmanis, A. C. Hurt, J. W. Yewdell, S. J. Kent, and A. K. Wheatley. 2018. Subdominance and poor intrinsic immunogenicity limit humoral immunity targeting influenza HA-stem. Journal of Clinical Investigation.

29. Baldrick, P., D. Richardson, G. Elliott, and A. W. Wheeler. 2002. Safety evaluation of monophosphoryl lipid A (MPL): An immunostimulatory adjuvant. Regul Toxicol Pharm 35: 398–413.

30. Jack, C., A. Adwani, and H. Krishnan. 2005. Tattoo pigment in an axillary lymph node simulating metastatic malignant melanoma. Int Seminars Surg Oncol 2: 28.

31. Litton, T. P., and S. V. Ghate. 2020. Tattoo pigment mimicking axillary lymph node calcifications on mammography. Radiology Case Reports 15: 1194–1196.

32. Matsika, A., B. Srinivasan, J. M. Gray, and C. R. Galbraith. 2013. Tattoo pigment in axillary lymph node mimicking calcification of breast cancer. Bmj Case Reports 2013: bcr2013200284–bcr2013200284.

33. Martin, J. T., C. A. Cottrell, A. Antanasijevic, D. G. Carnathan, B. J. Cossette, C. A. Enemuo, E. H. Gebru, Y. Choe, F. Viviano, S. Fischinger, T. Tokatlian, K. M. Cirelli, G. Ueda, J. Copps, T. Schiffner, S. Menis, G. Alter, W. R. Schief, S. Crotty, N. P. King, D. Baker, G. Silvestri, A. B. Ward, and D. J. Irvine. 2020. Targeting HIV Env immunogens to B cell follicles in nonhuman primates through immune complex or protein nanoparticle formulations. Npj Vaccines 5: 72.

34. Shukla, G. S., W. C. Olson, S. C. Pero, Y. Sun, C. L. Carman, C. L. Slingluff, and D. N. Krag. 2017. Vaccine-draining lymph nodes of cancer patients for generating anti-cancer antibodies. J Transl Med 15: 180.

35. Pero, S. C., Y.-J. Sun, G. S. Shukla, C. L. Carman, C. C. Krag, C. Teuscher, D. N. Krementsov, and D. N. Krag. 2017. Vaccine draining lymph nodes are a source of antigenspecific B cells. Vaccine 35: 1259–1265.

36. Shukla, G. S., Y.-J. Sun, S. C. Pero, G. S. Sholler, and D. N. Krag. 2018. Immunization with tumor neoantigens displayed on T7 phage nanoparticles elicits plasma antibody and vaccine-draining lymph node B cell responses. J Immunol Methods 460: 51–62.

37. Pauthner, M., C. Havenar-Daughton, D. Sok, J. P. Nkolola, R. Bastidas, A. V. Boopathy, D. G. Carnathan, A. Chandrashekar, K. M. Cirelli, C. A. Cottrell, A. M. Eroshkin, J. Guenaga, K. Kaushik, D. W. Kulp, J. Liu, L. E. McCoy, A. L. Oom, G. Ozorowski, K. W. Post, S. K. Sharma, J. M. Steichen, S. W. de Taeye, T. Tokatlian, A. T. de la Peña, S. T. Butera, C. C. LaBranche, D. C. Montefiori, G. Silvestri, I. A. Wilson, D. J. Irvine, R. W. Sanders, W. R. Schief, A. B. Ward, R. T. Wyatt, D. H. Barouch, S. Crotty, and D. R. Burton. 2017. Elicitation of robust tier 2 neutralizing antibody responses in nonhuman primates by HIV envelope trimer immunization using optimized approaches. Immunity 46: 1073–1088.e6.

38. Elmore, S. A. 2006. Histopathology of the lymph nodes. Toxicol Pathol 34: 425–454.

39. Boes, K. M., and A. C. Durham. 2017. Pathologic basis of veterinary disease. Sect Ii Pathology Organ Syst 724–804.e2.

40. Listinsky, C. M. 1988. Common reactive erythrophagocytosis in axillary lymph nodes. Am J Clin Pathol 90: 189–192.

41. Hewitt, R. J., C. Wright, D. Adeboyeku, D. Ornadel, M. Berry, M. Wickremasinghe, A. Wright, A. Sykes, and O. M. Kon. 2013. Primary nodal anthracosis identified by EBUS-TBNA as a cause of FDG PET/CT positive mediastinal lymphadenopathy. Respir Medicine Case Reports 10: 48–52.

42. Ringe, R. P., R. W. Sanders, A. Yasmeen, H. J. Kim, J. H. Lee, A. Cupo, J. Korzun, R. Derking, T. van Montfort, J.-P. Julien, I. A. Wilson, P. J. Klasse, A. B. Ward, and J. P. Moore. 2013. Cleavage strongly influences whether soluble HIV-1 envelope glycoprotein trimers adopt a native-like conformation. Proc National Acad Sci 110: 18256–18261.

43. Sliepen, K., B. W. Han, I. Bontjer, P. Mooij, F. Garces, A.-J. Behrens, K. Rantalainen, S. Kumar, A. Sarkar, P. J. M. Brouwer, Y. Hua, M. Tolazzi, E. Schermer, J. L. Torres, G. Ozorowski, P. van der Woude, A. T. de la Peña, M. J. van Breemen, J. M. Camacho-Sánchez, J. A. Burger, M. Medina-Ramírez, N. González, J. Alcami, C. LaBranche, G. Scarlatti, M. J. van Gils, M. Crispin, D. C. Montefiori, A. B. Ward, G. Koopman, J. P. Moore, R. J. Shattock, W. M. Bogers, I. A. Wilson, and R. W. Sanders. 2019. Structure and immunogenicity of a stabilized HIV-1 envelope trimer based on a group-M consensus sequence. Nat Commun 10: 2355.

44. Whittle, J. R. R., A. K. Wheatley, L. Wu, D. Lingwood, M. Kanekiyo, S. S. Ma, S. R. Narpala, H. M. Yassine, G. M. Frank, J. W. Yewdell, J. E. Ledgerwood, C.-J. Wei, A. B. McDermott, B. S. Graham, R. A. Koup, and G. J. Nabel. 2014. Flow cytometry reveals that H5N1 vaccination elicits cross-reactive stem-directed antibodies from multiple Ig heavy-chain lineages. J Virol 88: 4047–4057.

45. Juno, J. A., H.-X. Tan, W. S. Lee, A. Reynaldi, H. G. Kelly, K. Wragg, R. Esterbauer, H. E. Kent, C. J. Batten, F. L. Mordant, N. A. Gherardin, P. Pymm, M. H. Dietrich, N. E. Scott, W.-H. Tham, D. I. Godfrey, K. Subbarao, M. P. Davenport, S. J. Kent, and A. K. Wheatley. 2020. Humoral and circulating follicular helper T cell responses in recovered patients with COVID-19. Nat Med 1–7.

46. Schindelin, J., I. Arganda-Carreras, E. Frise, V. Kaynig, M. Longair, T. Pietzsch, S. Preibisch, C. Rueden, S. Saalfeld, B. Schmid, J.-Y. Tinevez, D. White, V. Hartenstein, K. Eliceiri, P. Tomancak, and A. Cardona. 2012. Fiji: an open-source platform for biological-image analysis. Nature Methods 9: 676.

47. Schneider, C. A., W. S. Rasband, and K. W. Eliceiri. 2012. NIH Image to ImageJ: 25 years of image analysis. Nature methods 9: 671–5.

